# Increased response to immune checkpoint inhibitors with dietary methionine restriction

**DOI:** 10.1101/2023.04.05.535695

**Authors:** Lauren C. Morehead, Sarita Garg, Katherine F. Wallis, Eric R. Siegel, Alan J. Tackett, Isabelle R. Miousse

## Abstract

Dietary methionine restriction, defined as reduction of methionine intake by around 80%, reproducibly decreases tumor growth and synergizes with cancer therapies. Here, we combined dietary methionine restriction with immune checkpoint inhibitors in a model of colon adenocarcinoma. *In vitro*, we observed that methionine restriction increased the expression of MHC-I and PD-L1 in both mouse and human colorectal cancer cells. We also saw an increase in the gene expression of STING, a known inducer of type I interferon signaling. Inhibition of the cGAS-STING pathway, pharmacologically or with siRNA, blunted the increase in MHC-I and PD-L1 surface and gene expression following methionine restriction. PD-L1 expression was also This indicated that the cGAS-STING pathway in particular, and interferon in general, is playing a role in the immune response to methionine restriction. We then combined dietary methionine restriction with immune checkpoint inhibitors targeted against CTLA-4 and PD-1 in a MC38 colorectal cancer tumor model in C57BL/6 mice. The combination treatment was five times more effective at reducing tumor size than immune checkpoint inhibition alone in males. We noted sex differences in the response to dietary methionine restriction for the MC38 tumor model in C57BL/6 mice. Finally, we observed an increase in PD-L1 protein expression in MC38 tumors from animals who were fed a methionine-restricted diet. Furthermore, the distribution of CD8 staining changed from mostly peripheric in the controls, to intratumoral in the methionine-restricted tumors. MHC-I, which has a high basal expression in MC38 cells, was highly expressed in all tumors. These results indicate that methionine restriction improves the response to immune checkpoint inhibitors in mice, and that this improvement is associated with the cGAS-STING pathway and interferon signaling.

## INTRODUCTION

Methionine is an essential amino acid. It is used not only for protein synthesis but also as a precursor for S-adenosylmethionine, the major cellular methyl donor. In healthy animals, reducing the food intake of methionine by about 80%, termed dietary methionine restriction, is associated with an increase in lifespan and an improvement of glucose and lipid regulation. In tumor-bearing animals, dietary methionine restriction decreases tumor growth and improves the response to radiation therapy (1,2). Other methionine-lowering approaches have also been shown to improve the response to chemotherapy (3–7) and targeted therapy (8). However, evidence showing that methionine restriction offers a benefit in combination with immune checkpoint inhibitors remains scarce.

Since their introduction in 2011, immune checkpoint inhibitors have rapidly changed the cancer field. Immune checkpoint inhibitors are antibodies that target immune molecules involved in dampening the anti-tumor immune response, the most common being CTLA-4, PD-1, and PD-L1. For specific types of tumors with a high tumor mutational burden, immune checkpoint inhibitors have increased response rates with remarkably few side effects. In metastatic colorectal cancer, anti-PD-1 therapy is approved as a first-line treatment for patients whose disease is characterized as either microsatellite instability-high or mismatch repair deficient (MSI-H/dMMR). This subtype represents 4-5% of patients with metastatic colorectal cancer (9). An anti-PD-1/anti-CTLA-4 combination currently in clinical trials has already generated encouraging results in patients with MSI-H-dMMR colorectal cancer, although randomized studies are still required (10). In MSI-H/dMMR rectal cancer, response rates up to 100% have been recorded (11). However, the MSI-H/dMMR subtype represents only a small fraction of all colorectal cancers. For the majority of patients, the efficacy of immune checkpoint inhibitors is limited by the poor immunogenicity of the tumor (12).

One important way by which tumors escape the immune system is by downregulating the expression of the major histocompatibility complex I (MHC-I) at their surface (13,14). MHC-I presents intracellular antigens to the immune system, including abnormal proteins produced by tumor cells. In human, the complex is formed of four domains; three domains from the α chain (encoded by HLA-A, HLA-B, and HLA-C in human) and one encoded by β_2_-microglobulin. The expression of the genes encoding the different components of MHC-I can be regulated by factors such as NF-κB binding to the enhancer A region and interferon regulatory factors binding to the interferon-stimulated response element (ISRE). Endogenous peptides emerging from the proteasome are then loaded on the MHC-I complex. In tumors with a high tumor mutational burden, many abnormal proteins are produced that are displayed to the immune system through MHC-I. The loaded complex is then imported into the endoplasmic reticulum by a transporter constituted of the TAP1 and TAP2 proteins for translocation to the cellular membrane. During its translocation to the membrane, other events such as folding, stabilization, and autophagy can influence the final abundance of MHC-I presented at the surface (15).

Several approaches have been tested to increase MHC-I expression on the surface of cancer cells in an effort to improve responsiveness to immune checkpoint inhibitors. Interventions that augment inflammation can lead to an upregulation of MHC-I. For example, TNFα increases the activity of NF-κB, leading to an increase in MHC-I (16). However, treatment with TNFα is associated with severe toxicity (17). Another way to increase MHC-I is through interferon signaling. Similarly to TNFα, the use of interferon therapy is limited by important side-effects (18).

Another marker that predicts the response to immune checkpoint inhibitor is PD-L1. PD-L1 expression by cancer cells downregulates the anti-tumor immune response by binding to the inhibitory PD-1 receptor on cytotoxic T-cells. Most immunotherapy regimens include an anti-PD-1/PD-L1 antibody, alone or in combination. High expression of PD-L1 by tumors indicate a reliance on this inhibitory mechanism and therefore represents one of the factors that affect the susceptibility to anti-PD-1 therapy (19). Several factors regulate PD-L1 expression, including interferon signaling.

Previous research has shown a beneficial effect from protein restriction in combination with immune checkpoint inhibitors (20). In addition, restricting specifically methionine in the diet has been reported to have effects on the immune system. In healthy animals, dietary methionine restriction delays the age-related decline in immunity. At 18 months of age, mice fed a methionine-restricted diet showed a pattern of T-cell subsets in the peripheral blood that was closer to the one of young animals. However, T cell differentiation is also reported to depend on methionine uptake (21). Because of this conflicting activity of methionine on the immune system, we investigated whether methionine restriction would have a positive or negative impact on the response to immune checkpoint inhibitors. We found that dietary methionine restriction upregulates MHC-I and PD-L1 in cancer cells, leads to an upregulation of the cGAS-STING immune pathway and interferon signaling, and improves the response to immune checkpoint blockade in mice.

## MATERIALS AND METHODS

### Cell culture

HT29 (CVCL_0320), MC38 (CVCL_B288) were obtained from ATCC and used between passages 4 and 12. B16F10 OVA cells were a gift from Dr. Alan Tackett. Cells were maintained in standard DMEM with L-glutamine, 4.5g/L glucose and sodium pyruvate (Corning, Corning NY) supplemented with 10% FBS (Corning) and 100 IU penicillin and streptomycin (ThermoFisher Scientific, Waltham MA). Methionine-free and control media were prepared from high glucose, no glutamine, no methionine, no cystine DMEM (ThermoFisher Scientific) supplemented with 10% dialyzed serum (12-14 kD)(R & D Systems, Minneapolis, MN), 100 IU penicillin and streptomycin (ThermoFisher Scientific), 4 mM L-glutamine, and 1 mM sodium pyruvate (ThermoFisher Scientific). L-cystine (Millipore-Sigma, Burlington, MA) was resuspended in 1M HCl and added to the cell media at a final concentration of 150 µM. For control medium, L-methionine (Millipore-Sigma) was resuspended in PBS and added to the cell media at a final concentration of 200 µM. For the methionine-restricted media, L-methionine was added at a final concentration of 5 µM. Unless otherwise indicated, assays were performed after 24 hours of treatment. For cGAS-STING inhibition, the STING inhibitor C-176 was used at a final concentration of 5 µM (Tocris Bioscience (Bio-Techne), Bristol, UK) and the cGAS inhibitor RU.521 was used at a concentration of 1 µM (MedChemExpress, Monmouth Junction, NJ) for 24 hours in both cases. The JAK inhibitor ruxolitinib was used at a final concentration of 4 µM (ThermoFisher Scientific).

### Gene expression analysis

To measure gene expression, we plated cells in a 6-well format in triplicates in standard DMEM media and let the cells attach. We then rinsed the cells in PBS and changed the cell culture media for treatment media as described above. RNA was extracted using QIAzol lysis reagent (QIAGEN, Germantown, MD) following the manufacturer’s instruction. RNA quality and quantity was assessed by spectrophotometry (Nanodrop, ThermoFisher Scientific). Reverse transcription was performed on 1 µg of purified RNA using the iScript Reverse Transcription Supermix (Bio-Rad, Hercules, CA). For each quantitative real-time PCR reaction, 20 ng of cDNA was used and amplified with the iTaq Universal SYBR Green Supermix (Bio-Rad), using technical duplicates. Fold changes in expression were calculated using the ΔΔC_t_ method with the mean of the technical duplicates for each biological triplicate (22). We used the internal control gene *Rplp0*. The primers used are listed in Supplementary Table S1.

### Protein expression analysis

To measure protein expression, we plated cells in 10 cm dishes and proceeded as described in “Gene expression analysis”. After incubation in the treatment media, cells were rinsed with PBS and detached with trypsin. The cells were spun at 1000g for 5 min at 4°C. The cell pellet was then rinsed once with PBS and resuspended in RIPA buffer supplemented with a protease inhibitor cocktail (Millipore-Sigma) and phosphatase inhibitor tablets (ThermoFisher Scientific) for 30 min at 4°C with agitation. The resulting lysate was spun at 10,000 g for 10 min at 4°C. The protein concentration was measured in the resulting supernatant using a BCA method (ThermoFisher Scientific). Immunoblotting was performed by applying the quantified protein lysate to a Bis-Tris gel (NuPAGE, ThermoFisher Scientific) and transferred unto a PVDF membrane. The membrane was probed with an anti-STING antibody (#50494, Cell Signaling Technology, Danvers, MA)

### Flow cytometry

To measure the expression of MHC-I and PD-L1 on the cell surface, we plated cells in a 6-well format in triplicates as described above. Cells were harvested with trypsin, rinsed, and 1 million cells were resuspended in PBS containing 1% fetal bovine serum (ThermoFisher Scientific). Fluorescently-labeled antibodies against MHC-I (Cat# 311406 and 114608), H-2K^B^ bound to SIINFEKL (Cat# 141604), and PD-L1 (Cat# 393610 and 124312) (Biolegend, San Diego, CA) were added at a final dilution of 1:100. Cells were counter-stained with DAPI at a final concentration of 1 ng/mL and analyzed on a BD LSR Fortessa instrument (BD Biosciences, Franklin Lakes, NJ). The resulting data was processed using the software FlowJo (BD Biosciences). For the assessment of reactive oxygen species, cells were treated with CTL and MR media as above. After 24h, plated cells were incubated with CellROX reagent (ThermoFisher) at a final concentration of 2 µM in PBS for 30 min at 37°C. Cells were then harvested and analyzed as above.

### Animal studies

This project was approved by the Institutional Animal Care and Use Committee at the University of Arkansas for Medical Sciences (IACUC #4119). C57BL/6 mice were purchased from Jackson Laboratories (Bar Harbor, ME, USA). Mice were kept under standardized conditions with controlled temperature and humidity and a 12 h light/dark cycle. Mice were 5 months of age at the beginning of the experiment and were fed an identical standard laboratory chow up to Day 5 of the experiment.

We injected mice subcutaneously with 4x10^5^ MC38 cells on the right hip (N=8 animals per group). After 5 days of incubation, we changed the diets to a methionine-restricted (MR) diet containing 0.12% methionine, or an otherwise identical control (CTL) diet containing 0.65% per weight methionine (Teklad Diets TD.190775 and TD.140520, Supplementary Table S2). We maintained these diets continuously until endpoint. On days 5, 8, 11, and 13, mice in 2 groups (CTL+ICI and MR+ICI) were injected intraperitoneally with monoclonal antibodies against PD-1 (RMP1-14, 250 μg per dose) and CLTA-4 (9D9, 100 μg per dose) (BioXCell, Lebanon, NH) in sterile PBS or a control PBS solution (25). We measured tumors daily from Day 5, and calculated tumor volumes using the formula V= (W^2^ x L)/2 as previously described (23,24). When tumors reached 500-600 mm^3^ or on day 35, we anesthetized the mice with isoflurane and retroorbital bleeding was performed. We allowed the collected blood samples to coagulate at room temperature in serum gel tubes (Sarstedt, Newton, NC), and we then centrifuged them at 10,000× g for 10 min at 4 °C. The serum was transferred to new tubes and flash-frozen until analysis. Tumors were harvested and fixed in 10% formalin for immunohistochemistry, or flash-frozen in liquid nitrogen. We repeated the experiment using only the diet under the same conditions to obtain material for IHC. Tumor volumes from these animals are presented along with data from the main experiment in figure 4.

### Immune cell analysis

At sacrifice, we collected whole blood from the retro orbital sinus of tumor-bearing animals. We measured blood parameters with a Vetscan HM5 Hematology Analyzer.

### Immunohistochemistry

To evaluate PD-L1, CD8, and MHC-1 expression in formalin-fixed paraffin-embedded tissue sections of mouse tumors, immunohistochemical staining was performed with conventional techniques using an avidin–biotin complex, diaminobenzidine chromogen, and hematoxylin counterstaining (26). Sections (4 µm thick) were deparaffinized, rehydrated and blocked for endogenous peroxidase, followed by incubation with rabbit anti-PD-L1 (64988, 1:100, Cell Signaling Technology), rat anti-CD8 (14-0808-80, 1:50, Invitrogen, Waltham, MA) and rabbit anti-MHC-1 (NBP3-09017, 1:50, Novus Biologicals, Centennial, CO) overnight at 4°C. This was followed by a 30-min incubation with the biotinylated goat anti-rabbit IgG (1:400, Vector Laboratories). The sections were subsequently incubated with avidin–biotin–peroxidase complex for 45 min (1:100, Vector Laboratories). Peroxidase binding was visualized using 0.5 mg/mL 3,3-diaminobenzidine tetrahydrochloride solution (Millipore-Sigma) then rinsed and counterstained with Harris’s hematoxylin, dehydrated, and cover slipped. The stained slides were scanned using an Aperio Scanner CS2 at 20×.

Stained slides were scored and were placed in categories including negative (-), weak (+), moderate (++) and strong (+++) based on their positive expression.

### Data analysis

The software GraphPad Prism 8.3.0 (GraphPad Software, San Diego, CA) was used to represent the data graphically and perform statistical analysis. Comparisons were made using an unpaired T test for pairwise comparisons, and 2-way ANOVA with Šidák’s multiple comparison test (CTL vs MR within each treatment group) for experiments with a 2x2 design. SAS v9.4 (The SAS Institute, Cary, NC) was used to analyze tumor volumes vs days post-injection, as follows: First, we log-transformed tumor volumes to stabilize variances and to facilitate statistical inference on ratio changes. If a mouse’s tumor volume was 0 mm^3^, then its log-volume was set equal to 0. Then we used the MIXED Procedure in SAS v9.4 to analyze the longitudinal log-volume data from males and females separately. For each sex, we modelled the data’s autocovariance structure with a heterogeneous 1^st^-order autoregressive model. We used the sex-specific models to estimate the mean log-volume with 90% confidence interval of each group on each post-injection day, and to compare the MRD+ICI and CTL+ICI groups at unadjusted α=0.05 for differences in their mean log-volumes on each day post-injection. Finally, we exponentiated the log-volume differences to obtain ratio changes in tumor volume, and we also exponentiated the group mean log-volumes with 90% confidence intervals to display them graphically as tumor-growth curves.

### Data Availability

No data was generated or analyzed in the reported study

### Results

#### Methionine restriction increases MHC-I expression in cancer cells

MHC-I and PD-L1 are two markers of the response to immune checkpoint inhibitors. To assess the possible interaction between methionine restriction and immune checkpoint blockade, we measured the effect of methionine restriction on the expression of these two markers *in vitro*. We treated HT29 human colon cancer cells for 24 hours in control DMEM media containing 200 µM methionine or in an otherwise identical media containing 5 µM methionine (methionine-restricted). We found that both the genes *HLAA* and *CD274* (encoding the A subtype of MHC-I heavy chain and PD-L1, respectively) were significantly elevated in methionine-restricted cells compared to control (1.7x, p<0.0001 and 1.6x, p=0.0018 respectively)(Fig.1 A and B). To elucidate whether this increase at the gene expression levels was functionally relevant, we measured protein surface expression by flow cytometry. We found that the surface detection of MHC-I and PD-L1 was indeed increased by methionine restriction (Fig. 1C and D). To assess antigen presentation, we used murine B16F10 OVA cells, which express the SIINFEKL ovalbumin peptide. In that cell line, methionine restriction also increased the abundance of MHC-I at the cell surface (Fig. 1E). Furthermore, SIINFEKL peptide presentation in the context of MHC-I at the surface of methionine-restricted cells was nearly double that of cells grown in control levels of methionine (Fig. 1E). These results indicates that methionine restriction increases MHC-I gene expression and that this increase in gene expression is associated with an increase in surface abundance in more than one cell line. Finally, methionine restriction also increases the display of antigens in the context of MHC-I on the cell surface in cancer cells.

**Figure 1.**
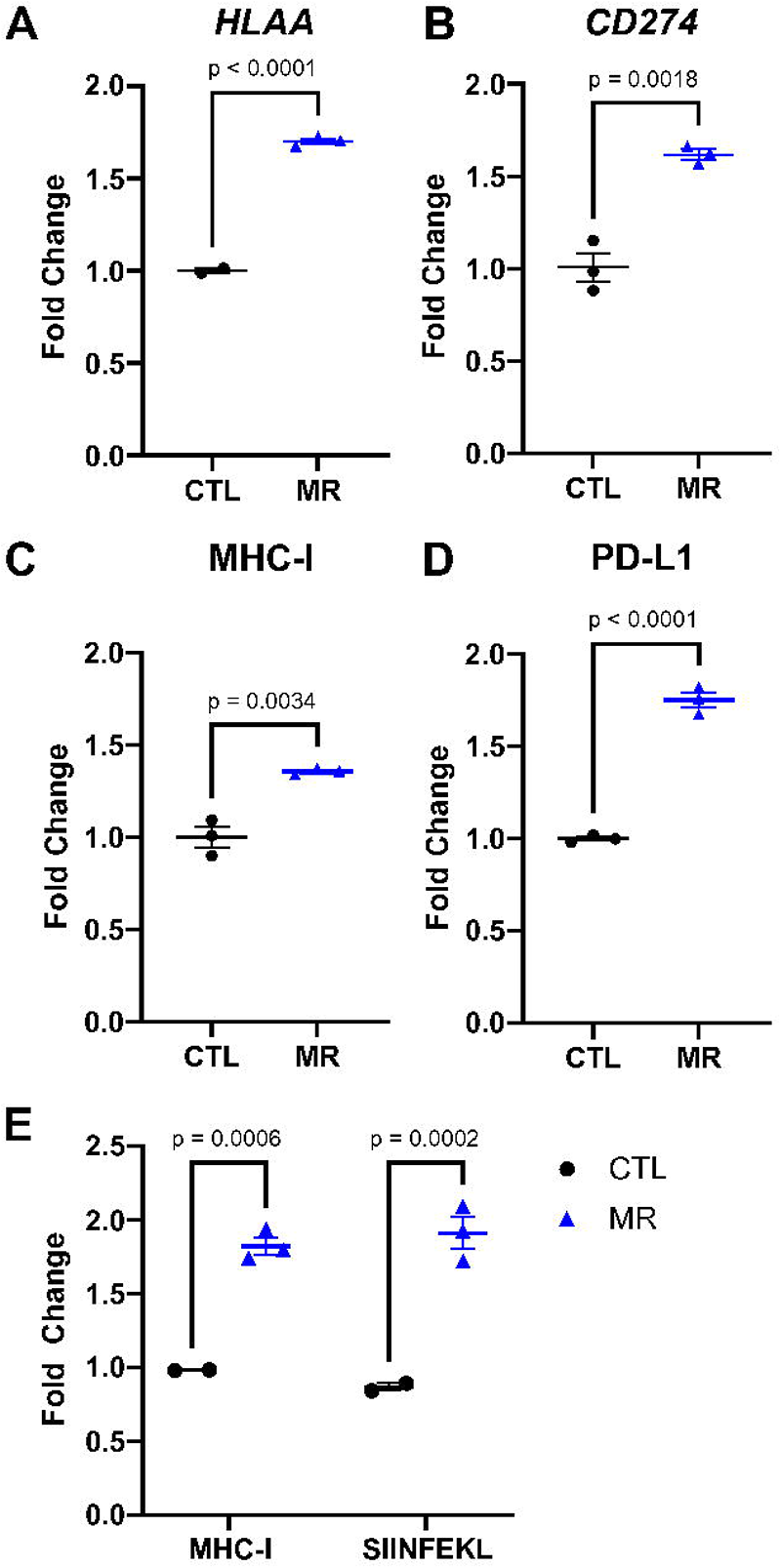
Methionine restriction increases MHC-I and PD-L1 expression. HT29 human colon cancer cells were grown for 24 hours in media containing 200 µM (CTL) or 5 µM (MR) methionine. Gene expression for *HLAA* (A) and *CD274* (B) increased with MR. Surface abundance measured by flow cytometry (C-D) also increased with MR. In B16-OVA cells, lowering methionine also increased MHC-I surface abundance and the presentation of the SIINFEKL peptide (E). T-test with mean and SEM shown.

#### The increase in MHC-I expression is STING-dependent

One of the triggers for MHC-I expression is the detection of cytosolic dsDNA through the cGAS-STING pathway. We hypothesized that inhibiting the cGAS-STING pathway would blunt the increase in MHC-I in response to low methionine. We observed that *STING* gene expression was elevated 1.6x in methionine-restricted HT29 cells (p=0.0012)(Fig. 2A). STING protein level was also elevated in methionine-restricted cells. We then treated cells with the STING inhibitor C-176 and analyzed *HLAA* gene expression. The increase in *HLAA* gene expression was blunted following STING inhibition, from 1.7x (p=0.0033) to 1.2x (p=0.0564)(Fig 2B). We found similar results when looking at MHC-I protein expression at the cell surface with flow cytometry (Fig 2C). The MHC-I increase in surface expression following methionine restriction was also blunted using the cGAS inhibitor RU-521 (Supp Fig S1). Treatment with C-176 had less clear effects on the *CD274* gene (PD-L1) expression. C-176 itself led to an increase in *CD274* expression, without further increase with the combination of C-176 and methionine-restriction. We therefore cannot definitely conclude from this result that STING expression is associated with the increase in *CD274* (Fig 2D). We also saw an increase in gene expression for *HLAB*, *HLAC*, as well as for interferon α and β (Supp Fig S1). These results suggest that the increase in MHC-I expression in colon cancer cells following methionine restriction is mediated at least in part through the cGAS-STING pathway.

**Figure 2.**
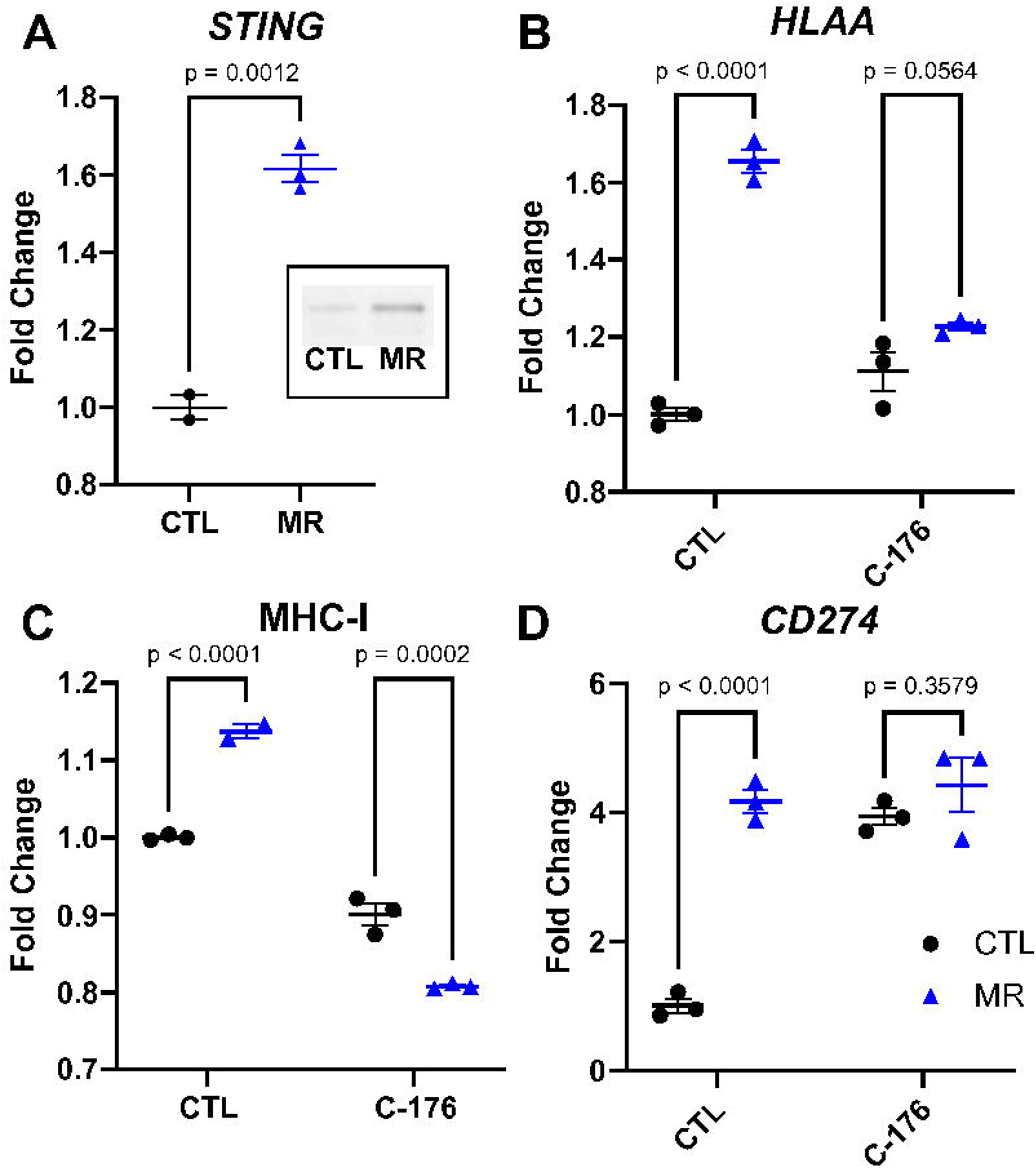
Inhibiting STING blunts the increase in MHC-I. *STING* was increased with MR at the gene and protein expression levels (A and insert, T-test with mean and SEM). Inhibiting *STING* with C-176 led to a reduction in the increase in *HLAA* gene expression (B) and in its cell surface representation (C). CD274 gene expression was increased by C-176 and did not increase further with MR (D). 2-way ANOVA with mean and SEM.

#### Dietary methionine restriction increases PD-L1 expression in the MC38 adenocarcinoma model cell line

After establishing that methionine restriction elevates MHC-I expression *in vitro*, we investigated if this increase translates into an improvement in the response to immune checkpoint inhibitors *in vivo*. MC38 murine colon adenocarcinoma cells are commonly used to model immune checkpoint inhibitors in human because they are syngeneic in C57BL/6 and can therefore form tumors in immunocompetent animals. Similarly to the human colon cancer cell line, methionine restriction led to an increase in the MHC-I component gene *H2Kb* in murine MC38 cells *in vitro* (Fig. 3A). However, when we analyzed the surface expression of MHC-I in MC38 cells, methionine restriction led to an unexpected decrease in MHC-I presence at the surface (Fig 3B). For PD-L1 on the other hand, both the gene expression and surface protein expression were significantly increased (Fig 3C and D). While STING signaling is dependent on type I interferon signaling, the regulation of PD-L1 is thought to be more closely related to type II interferon. We used the type I/II interferon signaling inhibitor Ruxolitinib. In this case, the increase in PD-L1 gene expression flowing methionine restriction was reduced by more than half. Ruxolitinib also blunted the increase in H2Kb gene expression (Supp Fig S2). Methionine restriction was also associated with an increase in various components of the interferon system (Supp Fig S2). These results indicate that interferon signaling is involved in the increase in PD-L1 gene expression in response to methionine restriction.

**Figure 3.**
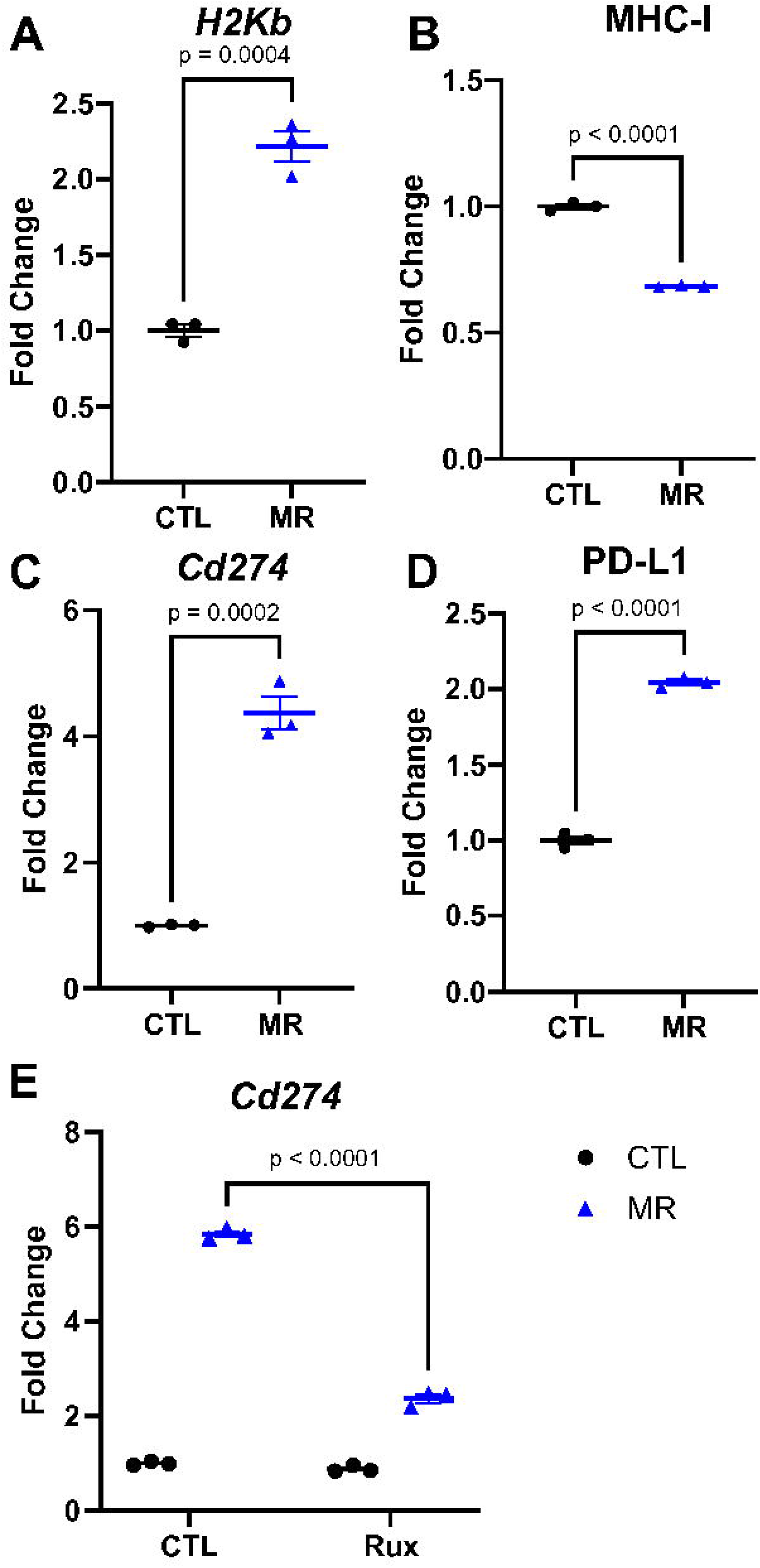
MHC-I and PD-L1 are altered by methionine restriction in a model cell line. Methionine restriction increases *H2Kb* gene expression *in vitro* in the cell line MC38 (A). The cell surface expression of MHC-I, however, decreased with methionine restriction (B). Gene expression for *Cd274* also increased with MR (C), with a parallel increase in surface protein abundance (D). T-test with mean and SEM. Inhibiting JAK (interferon signaling) with 4 uM ruxolitinib led to a blunting of the effect of MR on Cd274 gene expression (E). 2-way ANOVA with mean and SEM.

#### Dietary methionine restriction improves the response to immune checkpoint inhibitors *in vivo*

We injected MC38 cells subcutaneously on the hip of immunocompetent adult C57BL/6 mice and let the tumors develop. After 5 days, we changed the animal diets for a control amino acid defined diet containing 0.65% per weight methionine or an otherwise identical diet containing 0.12% methionine (18% of control). We maintained animals on these diets until endpoint. In females, treatment with the methionine-restricted diet alone did not lead to a significant reduction in tumor volume by day 15 after injection (10 days after diet initiation) (Fig 4A). In males on the other hand, the methionine-restricted diet led to a significant reduction in tumor volume (Fig 4B). Previous results had indicated a beneficial effect of methionine restriction in younger female mice (27), indicating that sexual maturity may influence the response (see discussion).

**Figure 4.**
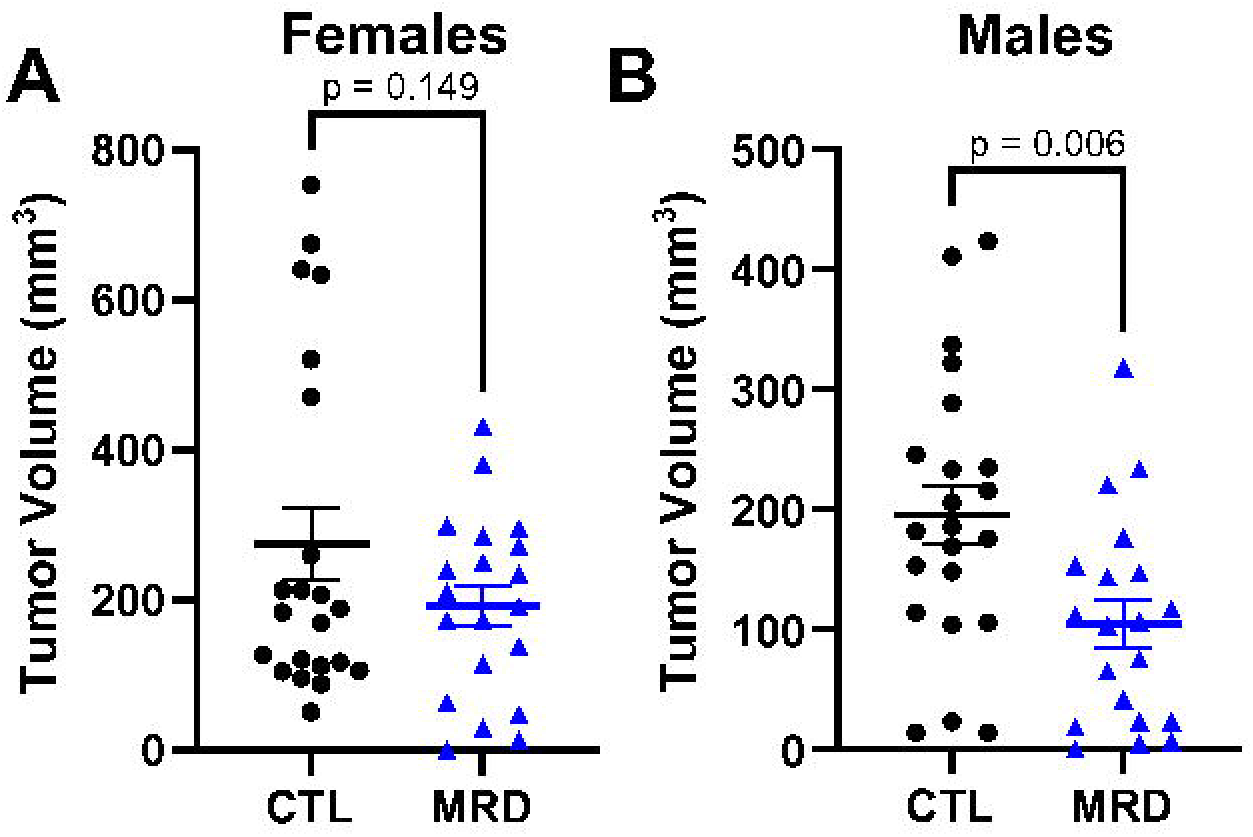
Dietary methionine restriction reduces tumor volumes in males. Immunocompetent females and males C57BL/6 mice were injected with tumor cells and five days later switched to a diet containing 0.65% methionine (CTL) or 0.12% methionine (MRD). Tumor volumes were calculated at day 15 after injection. Although we did not measure a difference in tumor volume in females (A), we observed a significant decrease in tumor volume at day 15 after injection in males (B). T-test with mean and SEM.

Based on these results, we assessed the effect of dietary methionine restriction on the response to immune checkpoint inhibitors. As before, we injected MC38 cells subcutaneously on the hip. Mice were 3 months of age at the beginning of the experiment. After five days, we initiated therapy with immune checkpoint inhibitors and the diet concomitantly. The immunotherapy regimen consisted of four intraperitoneal injections every three days of a combination of anti-PD-1 and anti-CTLA-4 monoclonal antibodies. Treatment with immune checkpoint inhibitors alone led to a 72-73% reduction in MC38 tumor volume in both females and males (Fig 5A). In females, we did not observe any improvement in the response to immune checkpoint inhibitors in animals consuming a methionine-restricted diet (Fig 5A). In males on the other hand, the combination treatment led to a further reduction in tumor volume, the combined effect being 5x greater than with immune checkpoint inhibitors alone (Fig. 5A and B, Supplementary Fig S3). As expected from a relatively mild methionine restriction, we did not note any significant difference in body weight between groups (Fig 5C, Supplementary Fig S3).

**Figure 5.**
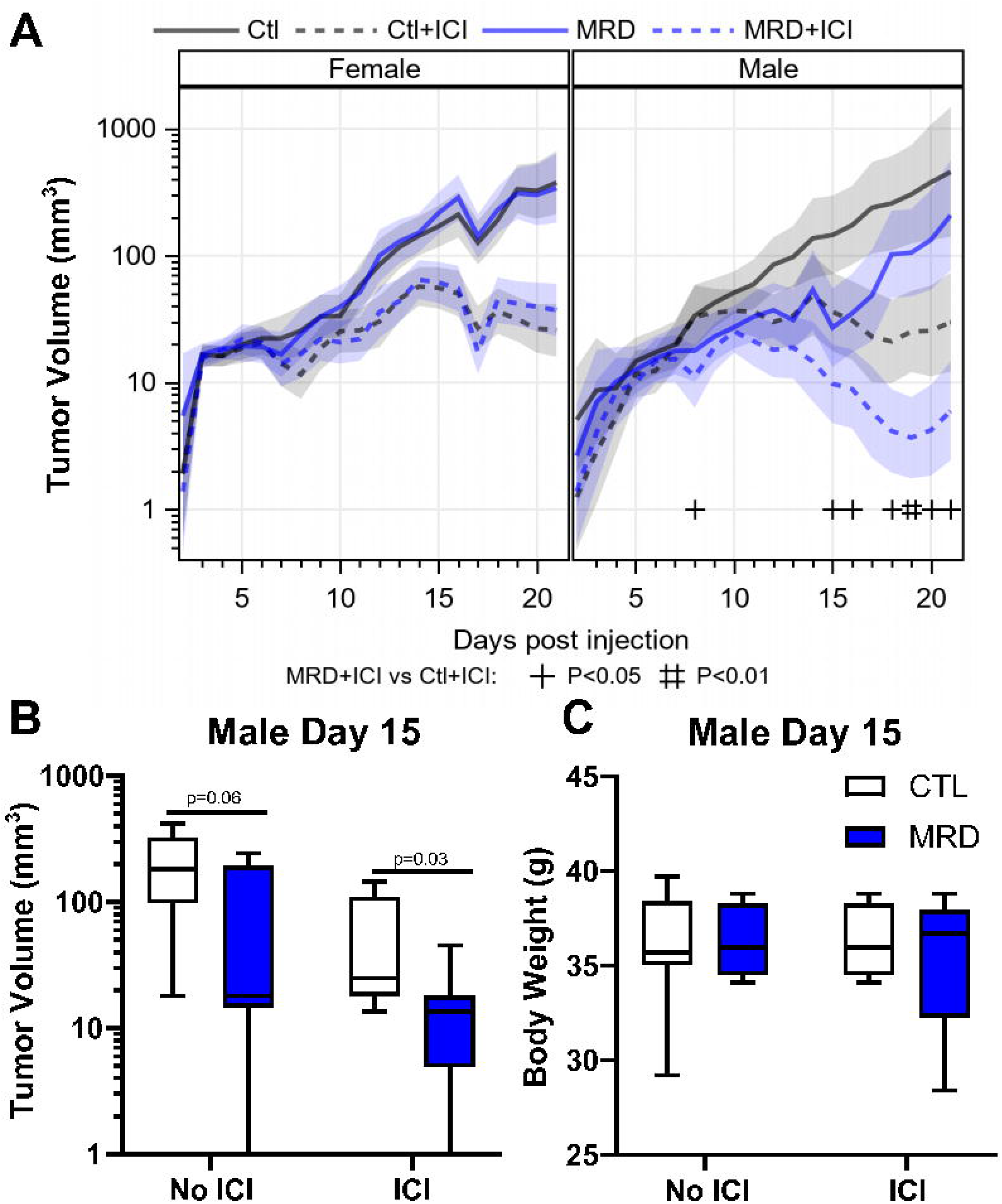
Dietary methionine restriction synergizes with immune checkpoint inhibitors in males. Upon combination of treatment with anti-CTLA-4 and anti-PD-1 with MRD, females showed no benefit from MRD (A). In males, there was a 5X reduction in tumor volume with immune checkpoint inhibitors (ICI) and MRD compared with ICI alone (B and C). There was no significant change in body weight measured after 10 days of MRD (C).

#### Dietary methionine restriction increases PD-L1 expression and CD8 infiltration *in vivo*

To confirm that the improvement in tumor control observed with the methionine-restricted diet is also associated with an increase in MHC-I and PD-L1, we performed immunohistochemistry. We injected animals with MC38 cell subcutaneously as before and initiated the diets after five days. We harvested then fixed the tumors when they reached 500-700 mm^3^. Tumor volumes in males and females were consistent with our previous observations. We did not measure any significant change in white blood cells or lymphocytes associated with the diets in animals of either sex (Supplementary Fig S4). There was a significant decrease in hemoglobin in the methionine-restricted group in both females and males, with a small but significant decrease in red blood cells in females. To our knowledge, hemoglobin values have not been previously reported in methionine restriction but are in line with data reported for complete methionine starvation in acute myeloid leukemia (28). We then performed immunohistochemistry on the tumor tissue. Our analysis did not identify any change in MHC-I staining in the sections from animals consuming a methionine-restricted diet (Table 1). However, we did observe an increase in PD-L1 staining associated with methionine restriction in males (Table 1 and Fig. 6). This is in good agreement with the increase in PD-L1 gene and protein expression observed *in vitro* in MC38 cells after methionine restriction. We did not identify any change in PD-L1 staining in tumors from female animals (Table S3). We also observed that staining for CD8 was mostly restricted to the periphery of tumors in the animals consuming a control diet (3 tumors out of 5). On the other hand, all the tumors in animals consuming a methionine restricted diet displayed CD8 infiltration (Table 1 and Fig. 6).

**Figure 6.**
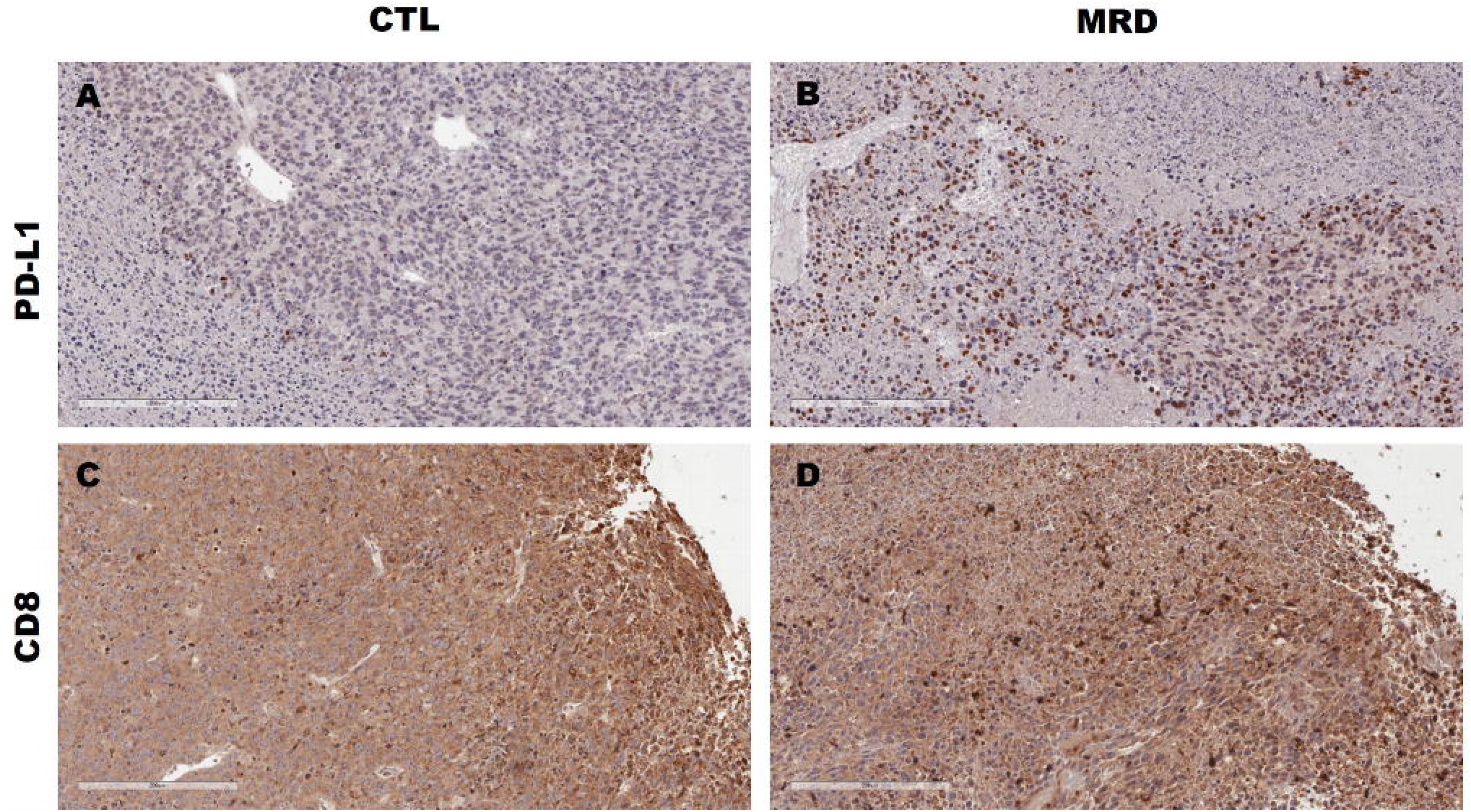
PD-L1 and CD8 in tumors from restricted animals. PD-L1 protein expression (brown) was higher in formalin-fixed tumor tissue from methionine-restricted animals (B) compared to control (A). For CD8, we observed a higher frequency of immune-excluded tumors in the control group (C) compared to the MRD group (D). Representative images at 20X are shown. The scale bar represents 200 µM.

**TABLE 1:**
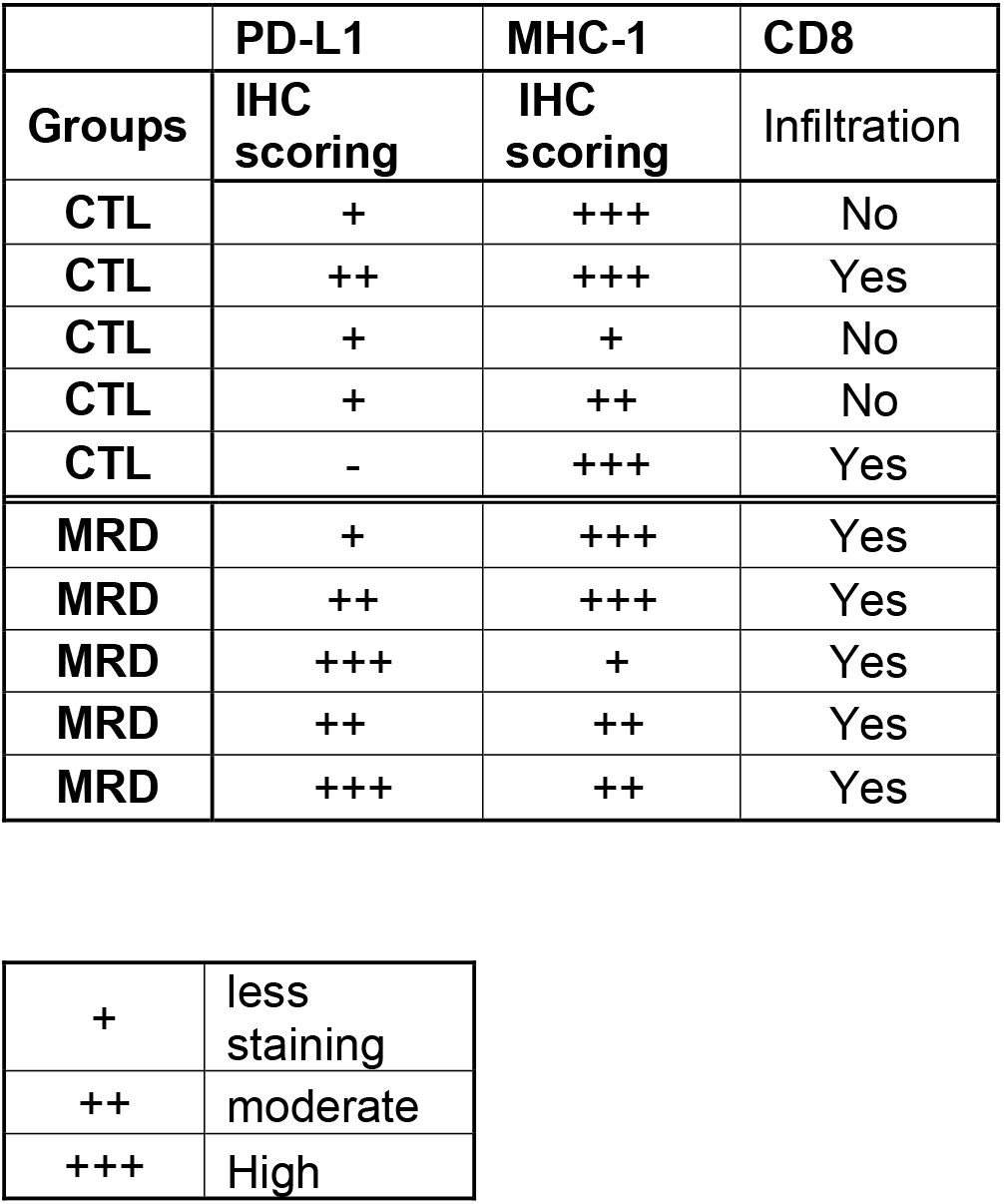
Effects of MRD on protein expression in MC38 tumors

|  | PD-L1 | MHC-1 | CD8 |
| --- | --- | --- | --- |
| Groups | IHC scoring | IHC scoring | Infiltration |
| CTL | + | +++ | No |
| CTL | ++ | +++ | Yes |
| CTL | + | + | No |
| CTL | + | ++ | No |
| CTL | - | +++ | Yes |
| MRD | + | +++ | Yes |
| MRD | ++ | +++ | Yes |
| MRD | +++ | + | Yes |
| MRD | ++ | ++ | Yes |
| MRD | +++ | ++ | Yes |

|  |  |
| --- | --- |
| + | less staining |
| ++ | moderate |
| +++ | High |

## DISCUSSION

The use of immune checkpoint inhibitors as a therapy for cancer has had a major impact on both survival and quality of life for patients. Immune checkpoint blockade has been approved in a subtype of metastatic colorectal cancer with high tumor mutational burden. However, its efficacy is limited in more common subtypes with lower mutational burden. We described here that dietary methionine restriction improves the response to immune checkpoint inhibitors in a model of colorectal cancer. Methionine restriction leads to an increase in the expression of PD-L1, whose interaction with PD-1 is the main target of immune checkpoint inhibitors. This increase was seen both *in vitro* and *in vivo*. We also found an increase in MHC-I, which displays antigens at the cell surface for surveying by the immune system, in the HT29 human colon cancer cell line.

The increase in MHC-I expression translated into an increased display of antigens on the cell surface, as attested by the increase in the presence of the SIINFEKL peptide conjugated to MHC-I at the surface of murine B16F10 OVA cells. Our results indicate that the increase in MHC-I expression is mediated at least in part through the cGAS-STING pathway. STING inhibition with C-176 blunted the increase in MHC-I after methionine restriction, both at the gene expression level and at the surface protein expression level. A similar activation of the cGAS-STING pathway has been described in response to a decrease in a different amino acid; arginine (29). Furthermore, we also found evidence of type I interferon signaling (interferon α and β) in response to methionine restriction that support our cGAS-STING observations. However, we cannot exclude the possibility that additional pathways may have a contribution.

Although the expression of PD-L1 was slightly affected by blocking the cGAS-STING pathway, PD-L1 is more closely associated with type II interferon (ɣ) signaling (30,31). In line with our results, recent evidence points to some degree of cross-talk between interferon ɣ and STING (32). We used the JAK inhibitor ruxolotinib to demonstrate that blocking both type I and type II interferon signaling resulted in a blunting of the PD-L1 response to methionine restriction. These results are also in good agreement with previous results showing an increased in interferon ɣ in methionine-restricted tumors (27). We believe that additional factors such as NRF2, already well described in the response to methionine restriction in cancer cells (33,34), may also contribute to the increase in PD-L1.

We then focused our attention to the interaction between immune checkpoint inhibitors and dietary methionine restriction in an animal model. Figure 4 shows the compiled tumor volume data at day 15 from three separate experiments. While males responded to the diet with a reduction in tumor volume as expected, females did not show a response. This was unexpected as previously published data indicated a response to methionine restriction in young C57BL/6 female mice injected with MC38 cells (27). Notably, Li and colleagues injected the cells in 7-week-old animals compared to 3-to 5-months-old animals in figure 4. This suggests that sexual maturity may influence the response to methionine restriction, especially in the context of the effects of sex on antitumor immune responses (reviewed in (35,36)). These differences may be particularly relevant to the increasing population of premenopausal women with early-onset colorectal cancer.

When we combined dietary methionine restriction with immune checkpoint inhibitors in the MC38 model, we observed a 5x improvement with the combination treatment compared to immune checkpoint inhibitors alone. This indicates that the changes that we observed *in vitro* may have a direct consequence *in vivo* on the response to therapy. However, there was no measurable benefits in females to the combination of methionine restriction and immune checkpoint inhibitors for MC38 tumors. MC38 is a cell line derived from a female animal, which may be expected to raise a weaker immune response in females. However, the similarity in response to ICIs between the males and females in the control diet argues against this. We previously reported a similar sex difference in the way the gut microbiome responds to dietary methionine restriction (37). There is accumulating evidence that the composition of the intestinal microbiome affects the response to immune checkpoint inhibitors (38–41). More experiments are required to identify what the contributions from the direct methionine levels are on tumors growth and responsiveness and indirect effects through the host such as glucose levels and microbiome composition.

Our results are supported by a previous study showing a benefit from combining protein restriction with immune checkpoint inhibitors in a model of renal cancer in females (20). In the same study, the authors also studied a methionine-restricted and cysteine-deficient diet in a model of prostate cancer in males. Methionine/cysteine restriction had a strong effect in delaying tumor growth. However, the additive effect between methionine/cysteine restriction and immune checkpoint blockade in the males was not statistically significant by the endpoint of 30 days. Several factors may have contributed to this result. The type of cancer was different – a slowly developing prostate cancer model versus colon adenocarcinoma here. Their diet also contained 0.17% L-methionine and 0% L-cysteine compared with 0.12% L-methionine and 0.35% L-cysteine in our work. Finally, Orillion and colleagues used anti-PD-1 therapy only, compared to the combination of anti-PD-1 and anti-CTLA-4 that we presented in this study. A recent report by Li and colleagues also supports the improvement of the response to a PD-1 only blockade in young female mice (27). However, Li and colleagues report a paradoxical decrease in PD-L1 with methionine restriction *in vitro*, which they attribute to RNA methylation. Unfortunately, the group did not report PD-L1 and MHC-I levels in the tumors. They however reported an increase in CD8 infiltration in MC38 tumors following methionine restriction that supports our own results.

In our experiment, we measured an increase in MHC-I gene expression in both the human and mouse colorectal cancer cell lines after methionine restriction *in vitro*. However, this increase did not lead to a corresponding increase in surface protein expression in the murine MC38 model *in vitro*. We also did not measure any change in MHC-I protein expression in tumors from methionine-restricted animals. MC38 has a high basal MHC-I expression, which may be of limited susceptibility to further protein-level increases at the membrane. We believe that the discrepancy between gene expression and membrane abundance is most likely due to an increase in endocytosis and/or in ubiquitylation of the protein. On the other hand, we did observe that PD-L1 surface expression was increased *in vitro* by methionine restriction, and that a larger proportion of methionine-restricted tumors had high PD-L1 expression than control tumors. The combination of strong basal MHC-I expression along with an improvement in PD-L1 expression resulting from the low methionine diet may have contributed to the more responsive phenotype that we measured. This also suggests that our study may underestimate the benefit of dietary methionine restriction for tumors that have a low basal expression of MHC-I.

Our results uncovered a previously undiscovered link between methionine restriction and the cGAS-STING pathway. Methionine restriction shares commonalities with caloric restriction and compounds referred to as “caloric restriction mimetics”. One of these compounds is the anti-diabetic drug metformin. Metformin has previously been shown to ameliorate the response to immune checkpoint inhibitors in animal models (42). Interestingly, metformin has also been reported to activate the cGAS-STING pathway in cancer cells (43). Our own results show that the caloric restriction mimetic and polyphenol resveratrol also increases genes via the cGAS-STING (submitted). As previously mentioned, decreases in the amino acid arginine also leads to the stimulation of the cGAS-STING pathway. The evidence suggests that this activation fits in a larger context of increased type I and type II interferon signaling. This suggests that other related compounds and interventions related to caloric restriction may similarly impact interferon signaling, and ultimately on the response to immune checkpoint inhibitors.

## ACKNOWLEDGEMENTS

The authors would like to thank the entire staff at the UAMS animal facility for facilitating this study, as well as Andrea Harris at the UAMS flow cytometry core. We are also grateful to Dr. Brian Koss for training and advice. This work was supported by the Winthrop P. Rockefeller Cancer Institute, the Arkansas Tobacco Settlement Commission, the National Center for Advancing Translational Sciences grant number 1KL2TR003108-01, and NIGMS grant P20 GM139768-01 (IRM) and R01CA236209-02 (AJT).

