## Supplementary figures and images for "Increased response to immune checkpoint inhibitors with dietary methionine restriction"

### Supplementary Figure 1

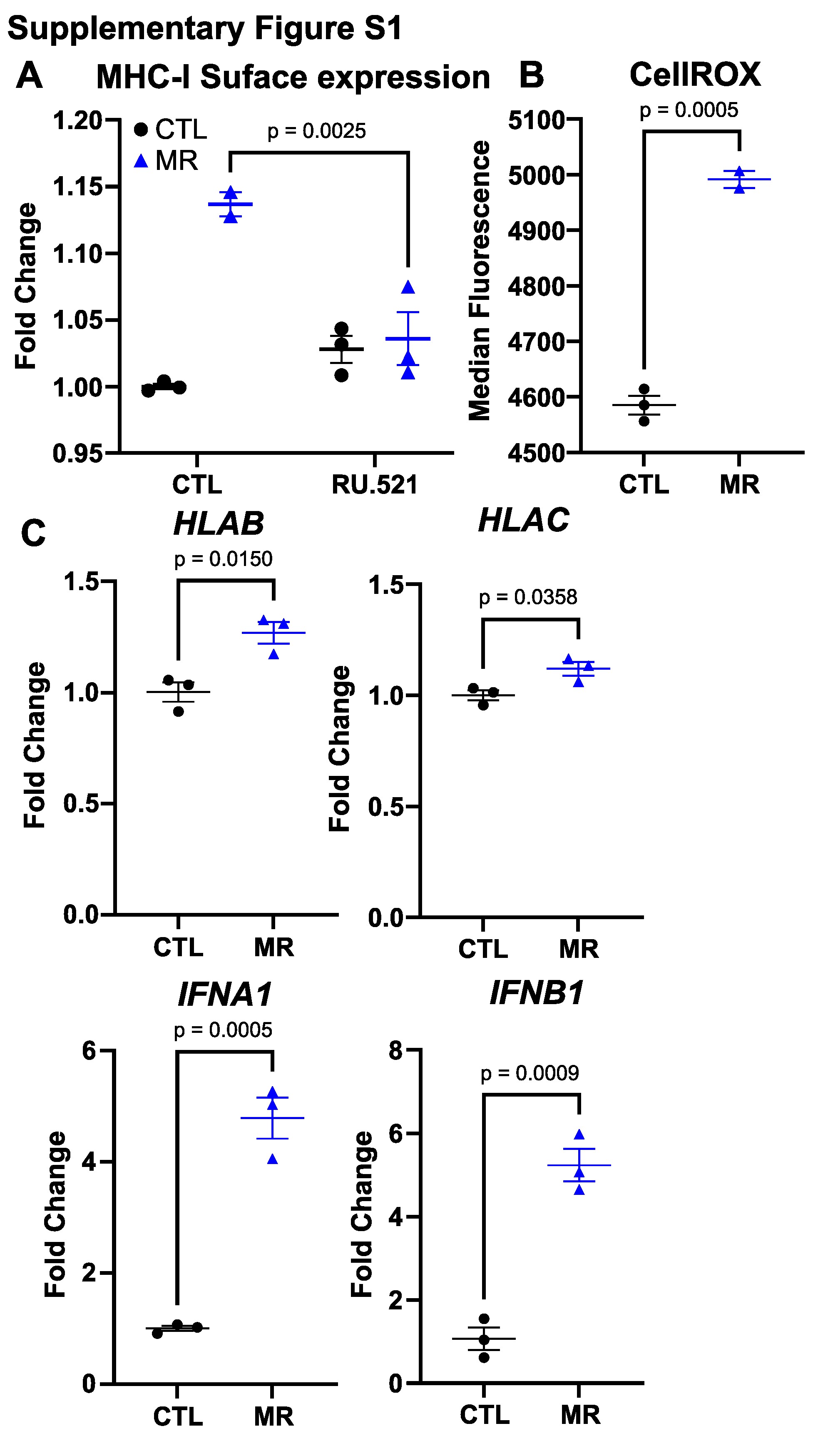

### Supplementary Figure 2

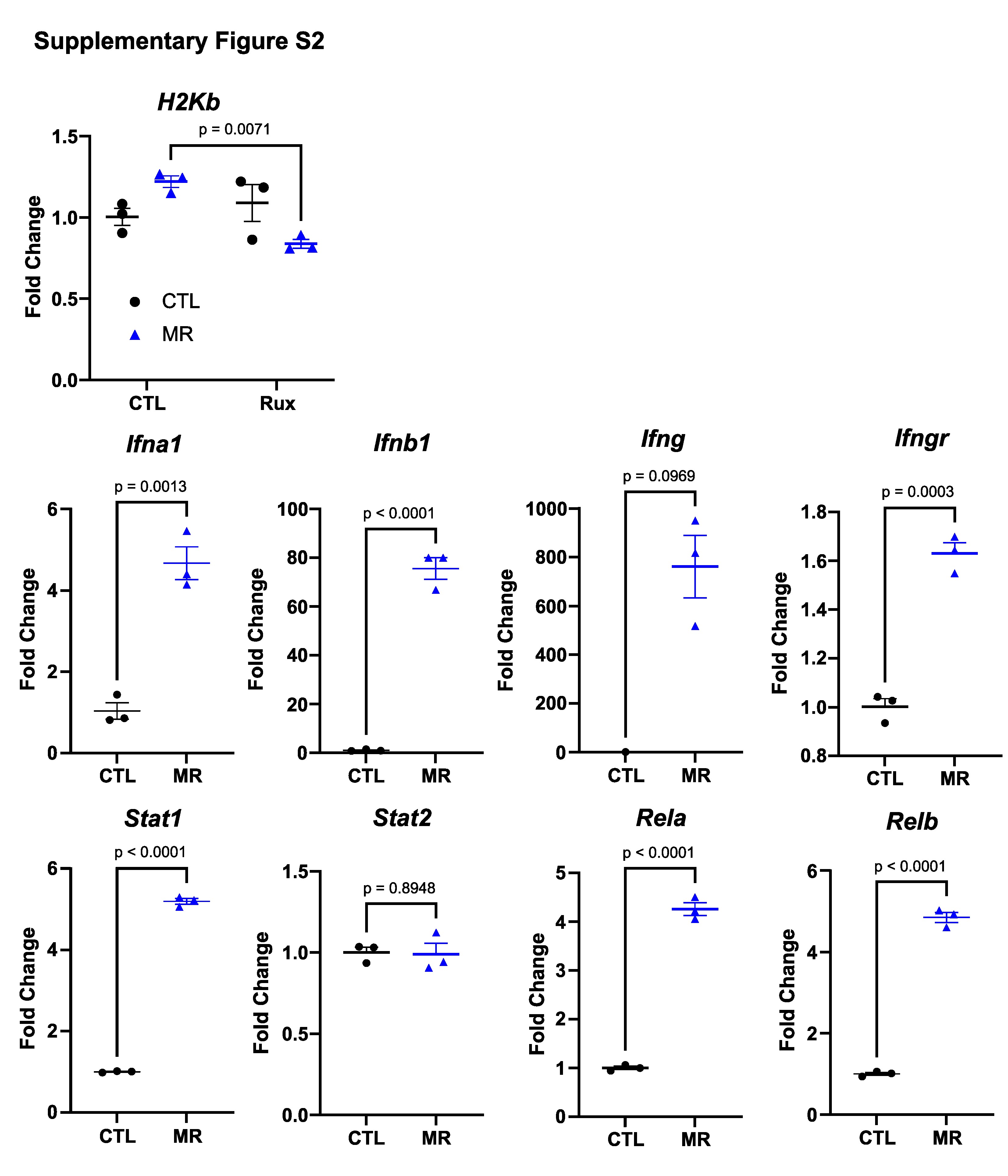

### Supplementary Figure 3

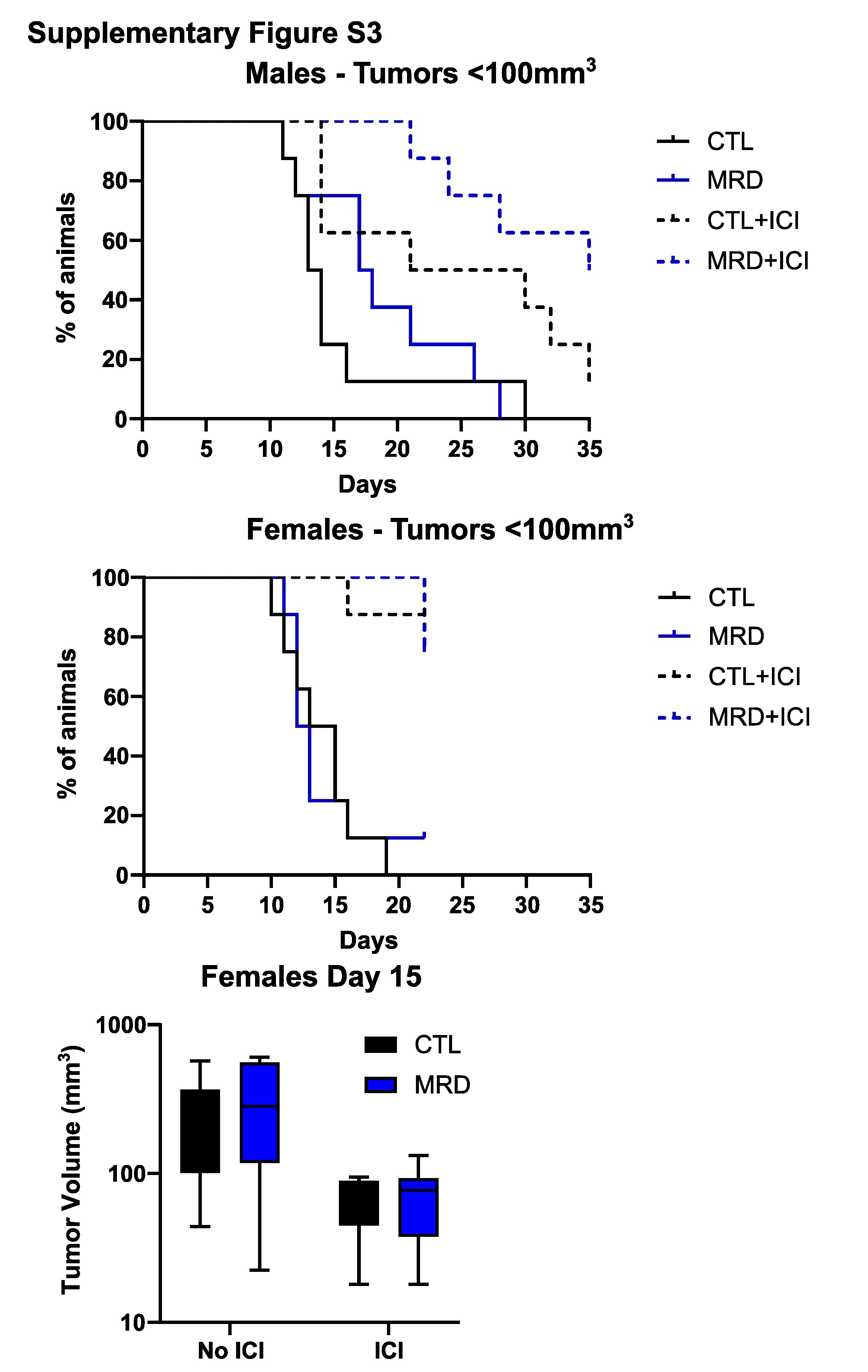

### Supplementary Figure 4

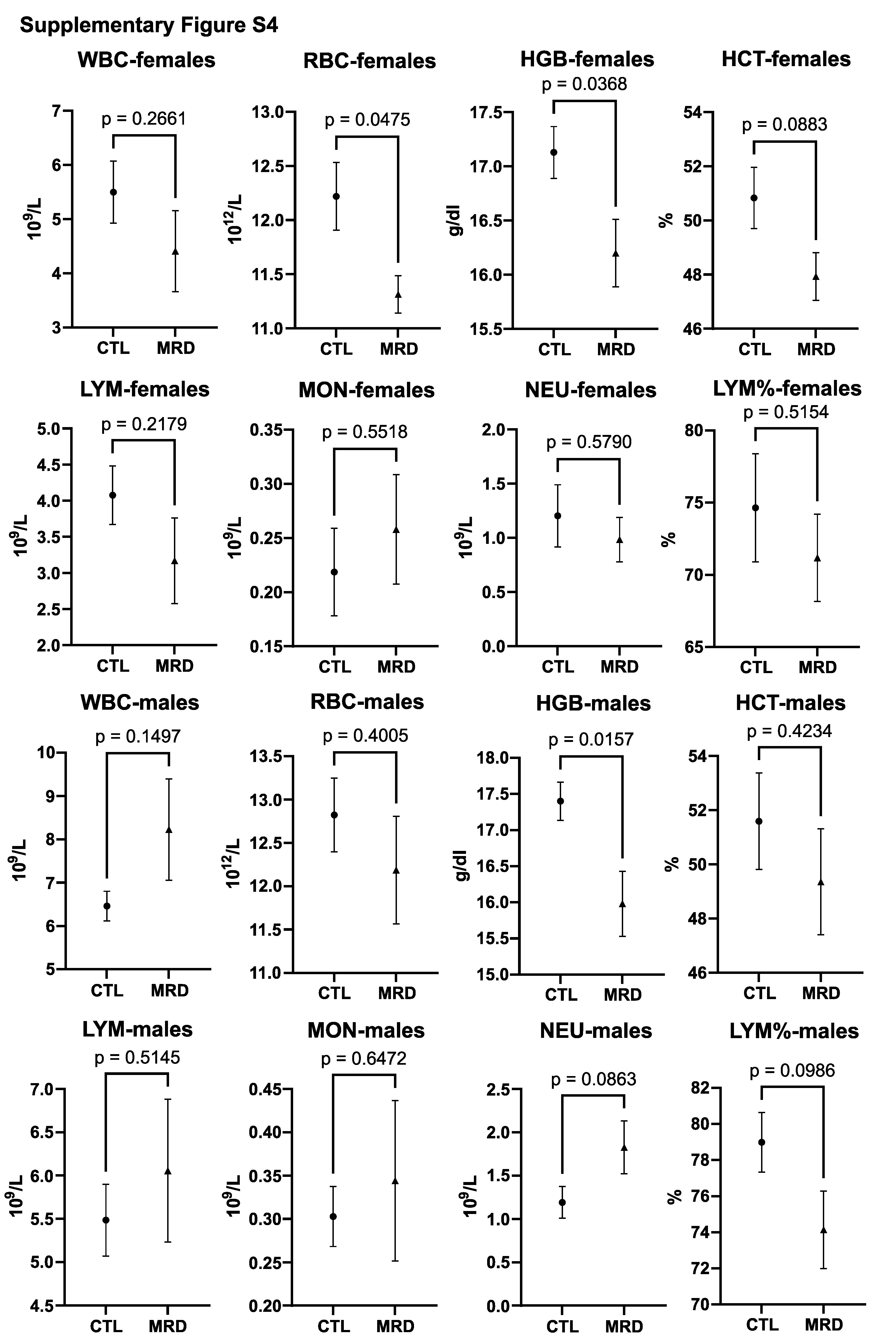
