## Supplementary Figure legend for "Increased response to immune checkpoint inhibitors with dietary methionine restriction"

Supplementary figure legends

**Supplementary Figure S1: Supplementary data for human HT29 cells *in vitro*.** The surface expression of MHC-I in response to MR was measured by flow cytometry in control HT29 cells and in cells treated with the cGAS inhibitor RU.512 at a final concentration of 1 µM (A). We measured an increase in oxidative stress as measured with the Cell ROX reporter in MR-treated cells (B). The median fluorescence is shown with SEM. The gene expression of *HLAB*, *HLAC*, *IFNA1* and *IFNB1* were significantly increased with MR (C). The fold change is shown with median and SEM.

**Supplementary figure S2: Supplementary data for murine MC38 cells *in vitro*.** The JAK inhibitor Ruxolitinib also blunted the increase in H2Kb gene expression. MR increased the gene expression for several interferon-related targets, except for *Stat2*. Interferon ɣ expression was undetectable in 2 out of 3 control samples. Fold change in expression is shown with the median and SEM.

**Supplementary figure S3. MC38 tumor volumes**. Kaplan Meier plots of MC38 tumor size representing the percentage of animals at the corresponding time point with tumors with a volume smaller than 100 mm^3^ in males and females. Box and whiskers plot of tumor volumes measured in females on day 15 post injection (min to max; box extends from the 25^th^ to 75^th^ percentiles).

**Supplementary figure S4. Blood parameters in tumor-bearing mice**. Blood parameters measured in animals bearing MC38 tumors at sacrifice. Mean with SEM is shown. WBC: white blood cells, RBC: red blood cells, HGB: hemoglobin, HCT: hematocrit, LYM: lymphocytes, MON: monocytes, NEU: neutrophils, LYM%: percentage of lymphocytes.
