## Supplementary Table 2 for "Increased response to immune checkpoint inhibitors with dietary methionine restriction"

**Table S1.** Nutritional characteristics of the methionine-adequate (TD.140520) and methionine-restricted (TD.190775) diets used in the study.

| **Formula** | **TD.140520**  **g/Kg** | **TD.190775**  **g/kg** |
| --- | --- | --- |
| Sucrose | 445.297 | 449.28 |
| Corn Starch | 198.783 | 200 |
| Corn Oil | 100 | 100 |
| Cellulose | 30 | 30 |
| Mineral Mix, AIN-76 (170915) | 35 | 35 |
| Calcium Phosphate, dibasic | 3 | 3 |
| L-Alanine | 3.5 | 3.5 |
| L-Arginine HCl | 12.1 | 12.1 |
| L-Asparagine | 6 | 6 |
| L-Aspartic Acid | 3.5 | 3.5 |
| L-Cystine | 3.5 | 3.5 |
| L-Glutamic Acid | 40 | 40 |
| Glycine | 23.3 | 23.3 |
| L-Histidine HCl, monohydrate | 4.5 | 4.5 |
| L-Isoleucine | 8.2 | 8.2 |
| L-Leucine | 11.1 | 11.1 |
| L-Lysine HCl | 18 | 18 |
| L-Methionine | 6.5 | 1.2 |
| L-Phenylalanine | 7.5 | 7.5 |
| L-Proline | 3.5 | 3.5 |
| L-Serine | 3.5 | 3.5 |
| L-Threonine | 8.2 | 8.2 |
| L-Tryptophan | 1.8 | 1.8 |
| L-Tyrosine | 5 | 5 |
| L-Valine | 8.2 | 8.2 |
| Vitamin Mix, Teklad (40060) | 10 | 10 |
| Ethoxyquin, antioxidant | 0.02 | 0.02 |
| % by weight | | |
| Protein | 15.3 | 14.9 |
| CHO | 63.3 | 63.8 |
| Fat | 10 | 10 |

*- Vitamin Mix, w/o choline, A, D, E; Vitamin E, DL-alpha tocopheryl acetate (500 IU/g; 0.242 g/kg); Vitamin A Palmitate (500,000 IU/g; 0.0396 g/kg); Vitamin D3, cholecalciferol (500,000 IU/g; 0.0044 g/kg).
