## Supplementary Table 3 for "Increased response to immune checkpoint inhibitors with dietary methionine restriction"

**Table S3. Effects of MR on PD-L1 expression in MC38 tumors in female mice**

|  | **PD-L1** |
| --- | --- |
| **Groups** | **IHC scoring** |
| **CTL** | + |
| **CTL** | + |
| **CTL** | + |
| **CTL** | + |
| **CTL** | + |
| **MRD** | + |
| **MRD** | + |
| **MRD** | + |
| **MRD** | ++ |
| + | less staining |
| ++ | moderate |
| +++ | High |
